# Minimum biomechanical energy expenditure predicts upper-limb motor strategies in individuals with limb loss

**DOI:** 10.1101/2025.02.27.640462

**Authors:** Christopher L. Hunt, Rebecca J. Greene, InHwa Lee, Rahul R. Kaliki, Nitish V. Thakor

## Abstract

Traditional models of upper-limb motion represent observed motor behaviors as the solution to an optimization problem defined over a cost function. However, these traditional formulations are computationally expensive and it is unclear if they extend to individuals with non-standard anatomy (such as those with upper-limb loss). *Goal:* We propose an optimal path planning framework that leverages musculoskeletal modeling to generate motor strategies during unconstrained, upper-limb movement. *Methods:* We validate this framework against upper-limb trajectories measured from a 3D target acquisition task and compare performance against multiple models of upper-limb motion previously presented in literature. *Results:* When compared to measured upper-limb trajectories, the proposed method generates upper-limb paths with significantly less geometric error than alternative methods (p < 0.001). *Significance:* Our approach provides a method for upper-limb motion planning that is easily adaptable to non-standard anatomies and computationally efficient enough for prosthesis control applications. *Conclusions:* The proposed path planning framework provides accurate motor strategy prediction for individuals both with and without upper-limb loss.

## I. Introduction

UPPER-limb human motion is remarkably consistent across individuals with several kinematic features being invariant to changes in movement size, speed, load, or direction [1], [2], [3], [4]. This regularity of features suggests that despite the infinite number of possible motor strategies capable of executing a desired motor task, the central nervous system (CNS) selects a single motor strategy among these possibilities using a reliable and reproducible method. Literature has described this motor strategy selection as an optimal control problem, with an observed motor strategy assumed to be the solution that minimizes some performance criteria (or cost function) [5], [6]. Literature has supported this conceptual framework by showing that the CNS does indeed evaluate and compare motor strategies based on their possible contributions to the current motor task [7], [8]. However, the workspace wherein this motor optimization occurs is an area of much debate. From inception to execution, a motor command undergoes several stages of transformation, all of which contain plausible criteria for optimization (Fig. 1).

**Fig. 1.**
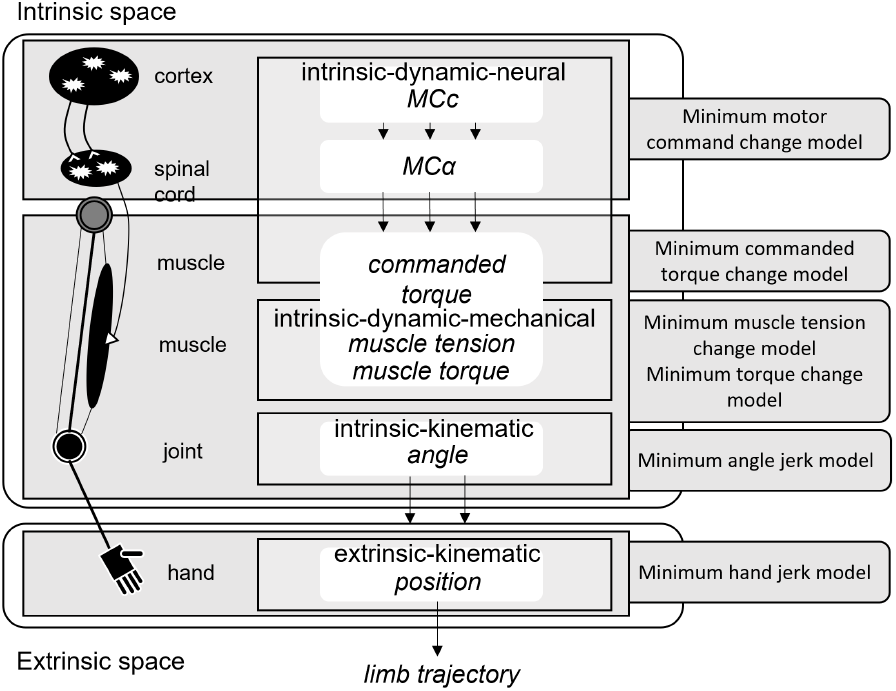
Conceptual workspaces for optimal trajectory formation. A motor command originates from the cortex (MCc) and is transmitted through the spinal cord via ***α***-motoneurons (MC***α***). These ***α***-motoneurons innervate musculature and generate tensions in the muscles they interface when activated. These tensions result in torques applied across the joints of the upper-limb, resulting in movement of the hand.

Several possible cost function models have been previously described, categorized into one of three broad categories: kinematic, dynamic, and energetic. Kinematic models directly optimize the geometry of a motor strategy and include criteria such as minimum hand jerk [9], minimum angle jerk [10], minimum angle acceleration [11], and minimum geodesic [12]. Dynamic models extend the optimization problem to consider the dynamics of the manipulators of the motor system (e.g. the upper-limb) and include criteria such as minimum torque [13] and minimum torque change [14], [15]. Energetic models further extend the optimization problem by considering the mechanical, neural, and biophysical energy expended to generate the observed motor strategy. Energetic models include criteria such as minimum absolute work [16], minimum effort [17], [18], and minimum metabolic energy expenditure [19], [20]. Of course, models may belong to multiple categories, including criteria like minimum endpoint variance [21]. Furthermore, while these models were originally proposed as alternatives to one another, recent evidence has shown that motor behavior is best described with a composite cost function, combining a weighted summation of the aforementioned costs [22], [23].

While this optimization framework does well to reproduce normative motor behavior, it is unclear how these methods generalize to individuals with partial upper-limb loss (ULL) [24]. During activities of daily living (ADLs), individuals with ULL have been shown to execute motor strategies with increased compensatory motion in the joints proximal to the amputation [25], [26], [27], decreased smoothness in joint actuation [28], and abnormal end-effector velocity profiles [29]. While unreliable prosthesis control schemes contribute to these observed motor differences [25], [26], literature has shown that the increased energy expenditure of the residual limb and prosthesis system influence the executed motor strategy as well [30].

In this study, we propose a motor prediction framework that aims to generate an individual’s upper-limb motor strategy given an initial and desired arm pose. The framework leverages an upper-limb musculoskeletal model to pre-compute a biomechanical cost space, over which we apply an optimal path planning algorithm to determine the lowest-cost path from the initial to desired arm pose.

## II Methods

### A. Experimental Protocol

This study was conducted in accordance with a protocol approved by the Johns Hopkins University School of Medicine Institutional Review Board. The participants consist of six individuals with intact limbs (IL) and one individual with partial upper-limb loss. IL participants include 4 males and 2 females that range in age from 25 to 29 years old and were right-handed dominant. The ULL participant was a 35-year-old male with bilateral upper-limb and lower-limb amputations required from meningitis complications. The ULL participant’s amputations include a transhumeral amputation on his left side, a wrist disarticulation on his right side, and partial foot amputations on both sides. The ULL participant was 11 years removed from his last upper-limb surgical revision and is right-hand dominant.

#### 1) Data Acquisition and Processing

Kinematics of the dominant limb were measured using a set of HTC VIVE trackers (HTC Corporation, New Taipei City, Taiwan). These kinematic trackers combine an infrared, optical tracking system with inertial measurement units to estimate the 3D position and orientation of each device. The devices were mounted on the shoulder, elbow, wrist, and hand of a participant’s dominant limb to record the upper-limb kinematics. Kinematic data was sampled at 90 Hz and was filtered using a 6-Hz second-order Butterworth low-pass filter in both the forward and backward directions, resulting in fourth-order filtering with zero-phase distortion [31].

#### 2) 3D Endpoint Acquisition Task

The primary motor task considered in this study is a 3D endpoint acquisition task (Fig. 2). Participants were seated, donning a virtual reality head-mounted display. Within the virtual reality environment, participants were prompted to place their dominant hand into a series of 3D locations, maintaining each posture for a brief moment.

**Fig. 2.**
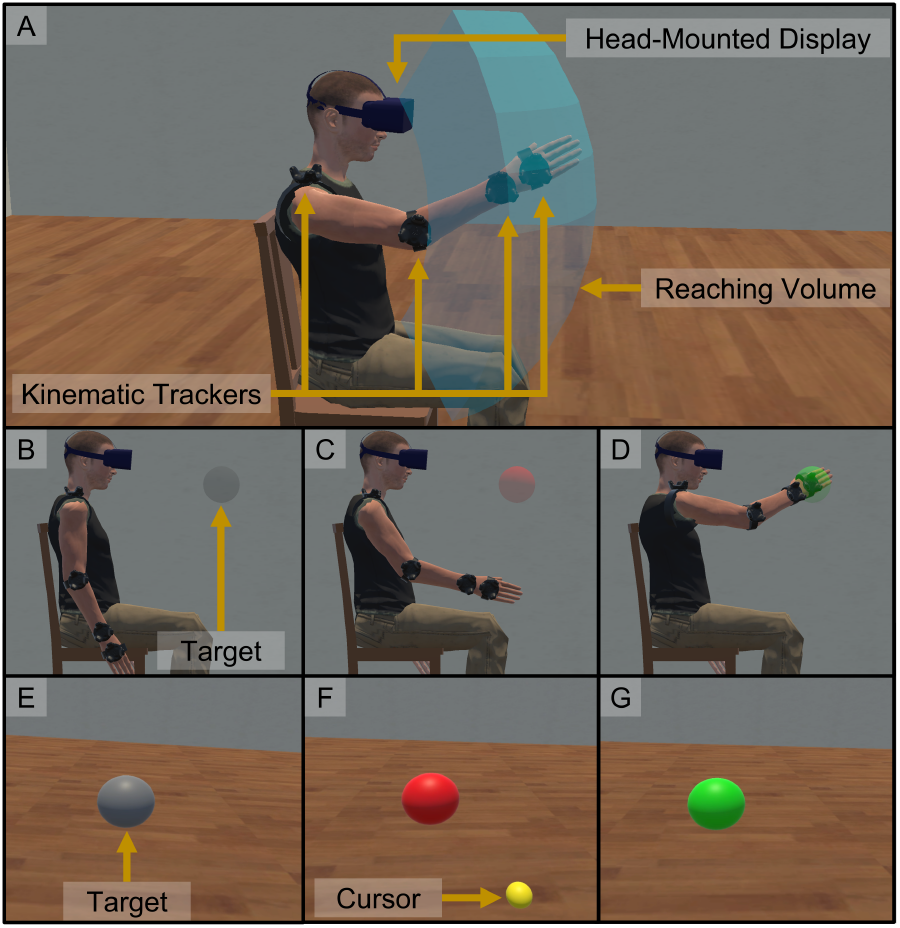
Virtual reality 3D endpoint acquisition task. (a) Participants were seated while wearing a virtual reality head-mounted display (HMD) and a series of kinematic trackers on their dominant limb. Participants were then tasked with placing their hand in a series of target 3D endpoint locations sampled from their reaching volume. (b) For a single trial, participants began with their dominant limb at their side with the target endpoint presented in the HMD as a grey sphere. (c) After 3 s, the target changed color to red, signifying that the participant should place their hand (represented as a yellow cursor) at the prompted location. (d) By holding the cursor at the target location for 1 s, the target sphere turned from red to green, signifying the end of the trial. A first-person view of a trial can be seen in subfigures (e – g).

For each participant, a total of 100 3D endpoint locations were prompted. Endpoint locations were uniformly sampled from the participants reaching volume, defined as the spherical subsection centered at the participant’s dominant shoulder and bounded by polar and azimuthal ranges: −45^°^≤ *α* ≤ 45^°^ and 45^°^≤ *β* ≤ 135^°^. The radial limits of the reaching volume were defined as 50% and 90% of the participant’s arm length, measured from the center of the shoulder to the distal tip of the middle finger (or residual limb).

For a single trial, participants began with their dominant limb at their side in a comfortable posture. A 3D endpoint target was then prompted as a grey sphere with a 10-cm diameter in the virtual environment placed within the participant’s reaching volume. After 3 s, the target sphere turned red, signaling that the participant was to place their hand (represented as a yellow sphere with a 5-cm diameter) at the prompted endpoint location. After the participant placed their hand within 10 cm of the target endpoint, the target sphere began changing color from red to green. The participant was then tasked with maintaining their hand within the 10-cm radius for 1 s continuously, until the target sphere turned completely green. Once the participant successfully positioned their hand within the 10-cm radius for 1 s, the trial would end and the participant would be prompted to return to the starting arm posture before beginning the next trial.

The ULL participant completed the 3D endpoint acquisition task twice, once while donning an upper-limb prosthesis using a bypass socket (i.e. *loaded*) and once while not wearing the prosthetic system (i.e. *unloaded*). The upper-limb prosthesis included a bebionic3 multiarticulated hand (Ottobock, Duderstadt, Germany) and a Motion Control Wrist Rotator (Fillauer LLC, Chattanooga, TN), weighing a total of 1.17 kg. After completing the unloaded condition, the participant was given the opportunity to rest before the loaded condition to counteract fatigue.

### B. Motor Path Optimization

#### 1) Musculoskeletal Model

We model a participant’s upper-limb using a musculoskeletal model (MSM) with 7 DoF: shoulder elevation plane, shoulder elevation magnitude, humeral rotation, elbow flexion, forearm rotation, wrist deviation, and wrist flexion [32]. The MSM contained *m* = 50 Hill-type actuators representing 32 muscles whose force dynamics follow Eq. 1:

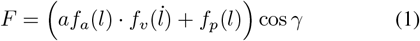

where *l* is the actuator length, *f*_*a*_ is the force-length relationship of the active muscle element, *f*_*p*_ is the force-length relationship of the passive tendon element, and *f*_*v*_ is the force-velocity relationship of the entire actuator. *a* is a scaling factor of the active element and *γ* is the pennation angle of the muscle fiber. A more detailed description of the dynamics of *f*_*a*_, *f*_*p*_, and *f*_*v*_ can be found in [33].

The total dynamics of the upper-limb can be defined as the sum of the actuator forces, ℱ= (*F*_1_, …, *F*_*m*_)^⊤^, acting across the lever arms, ℛ ∈ℝ^*m*×*n*^, of each respective DoF. From a Lagrangian mechanics perspective, the equations of motion can be defined as:

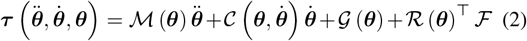

where ***τ*** = (*τ*_1_, …, *τ*_*n*_)^⊤^ is the set of generalized forces generated by the system and ***θ*** = (*θ*_1_, …, *θ*_*n*_)^⊤^ represents a set of upper-limb joint angles (i.e. an arm posture). While the subset (*θ*_1_, …, *θ*_7_)^⊤^ represents the 7 DoF of the MSM, the subset (*θ*_8_, …, *θ*_*n*_)^⊤^ represents the coupled joint angles that are not directly controllable in the MSM but do contribute to the total mechanical energy of the system. (e.g. sternoclavicular rotation, acromioclavicular rotation). ℳ, 𝒞 , and 𝒢 represent the inertia matrix, Coriolis/centrifugal matrix, and gravitational vector, respectively.

#### 2) Path Cost Estimation

Given the dynamic system defined in Eq. 2, we define the optimal upper-limb path **Θ**^∗^ as the curve through ℝ ^7^ that minimizes the absolute sum of the generalized forces:

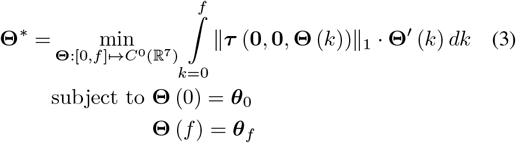

where ***θ***_0_ is the initial arm posture and ***θ***_*f*_ is the desired final arm posture. In this formulation, we have removed the dynamic contribution from the inertial and Coriolis/centrifugal terms found in Eq. 2. This is because literature suggests that the primary contributor to upper-limb dynamics generated during ADLs is the gravitational load, 𝒢 [34]. This implies that **Θ**^∗^ should depend primarily on the static forces generated by the gravitational load of the upper-limb. Eq. 3 captures this dependence by setting 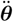 and 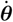 to **0** ∈ℝ^7^

Due to the non-convexity of Eq. 3, we solve for **Θ**^∗^ through numerical simulation of the MSM. For any arm posture, ***τ*** can be computed through an inverse dynamics simulation of the MSM in the OpenSim biomechanics software [35]. Normally, this method may be computationally expensive and not suited for a real-time application; however, because we have defined **Θ**^∗^ to not depend on the ℳ and 𝒞 components of the system dynamics, ***τ*** becomes time invariant, depending only on the current ***θ***. As such, we can pre-compute ***τ*** for a set of *N* arm postures, using this to construct a continuous cost function through interpolation. *N* = 20, 000 arm postures were selected via Halton sampling from the configuration space defined by the span of ***θ*** [36].

For each of the *N* sampled arm postures, we compute the generalized forces required to remain static in each arm posture for 1 s. ***τ*** is estimated to be the generalized forces at steady-state, generating a set of biomechanical costs, **c** = (*c*_1_, …, *c*_*N*_ )^⊤^ where *c*_*i*_ (*i* ∈ [1, *N* ]) is the *𝓁*_1_-norm of ***τ*** _*i*_. We then parameterize a function relating arm posture to biomechanical cost, using a radial basis interpolation with a multiquadric kernel [37]:

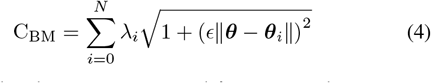

where *ϵ* is the shape parameter and *λ*_*i*_ is a weighting factor, computed through a least-squares optimization:

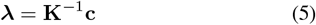

where ***λ*** = (*λ*_1_, …, *λ*_*N*_ )^⊤^ is the set of interpolant weighting factors, **c** is as previously defined, and **K** is the *N × N* , symmetric kernel matrix:

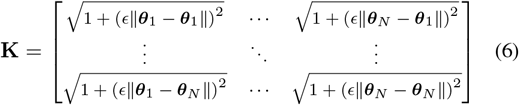

In this work, *ϵ* = 1 and the cost for each input ***θ*** was computed using the 50 nearest neighbors found within the set of *N* arm postures.

#### 3) Optimal Path Planning

Given C_BM_, the path from ***θ***_0_ to ***θ***_*f*_ is computed using a sampling-based planning algorithm: optimal transition-based rapidly-exploring random tree (or T-RRT*) [38]. Sampling-based planning algorithms work to find a path, ***π*** : [0, 1] → [***θ***_0_, ***θ***_*f*_ ], by sampling the configuration space (defined here as the span of ***θ***) and constructing a graph of nodes over which a feasible path is computed. In this work, T-RRT* was chosen for its following properties: (1) *asymptotic optimality* and (2) *accelerated convergence*. (1) T-RRT* is guaranteed to find ***π***^∗^ that minimizes the accumulated cost over C_BM_:

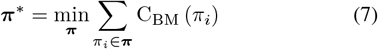

and (2) T-RRT* converges to find ***π***^∗^ more quickly than alternative planners. In this work, T-RRT* was implemented with 5000 sampling nodes. All other parameters were set to the values outlined in [38]. As ***π***^∗^ is not guaranteed to be smooth, **Θ**^∗^ was defined as the optimal path ***π***^∗^ interpolated via a cubic spline with a smoothing parameter of 0.9998 [39].

#### 4) Path Similarity Metric

Given **Θ**^∗^, we compute the resultant hand (or residual limb) path as:

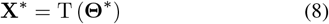

where T is the forward kinematic transformation of the MSM (derived from the OpenSim program) and **X**^∗^ is the optimal path in 3D Cartesian space, a sequence of points **x** = (*x*_1_, *x*_2_, *x*_3_)^⊤^.

To compute similarity between **X**^∗^ and a measured reference path **X**_ref_, we use the dynamic time warping algorithm (or DTW) [40], [41]. The DTW algorithm works to construct a *warping function ϕ* such that:

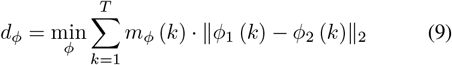

where *ϕ*_1_ and *ϕ*_2_ are the warped input paths and *m*_*ϕ*_ is a per-step weighting function. *d*_*ϕ*_ is the alignment distance, a global dissimilarity measure between two geometric paths. In this work, we use the DTW implementation found in [42] and report *d*_*ϕ*_ between a measured hand path and a computed optimal hand path as the primary performance metric.

#### 5) Comparative Models

The performance of the proposed biomechanical cost function, C_BM_, was compared to several models proposed in the literature: minimum hand jerk [9], minimum angle jerk [10], minimum torque change [14], [15], and minimum energy [16]. The formulations for these cost functions are summarized in Table I.

**TABLE I.**
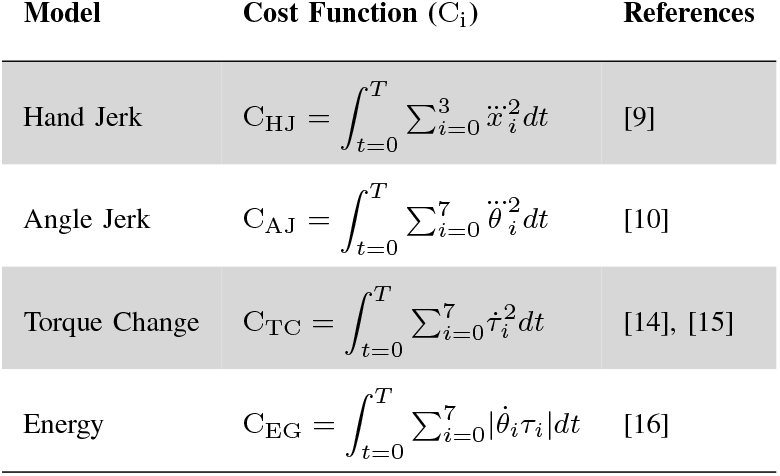
Comparative Cost Functions.

For models C_TC_ and C_EG_, the required dynamics calculations were computed by translating the MSM model to the MuJoCo physics engine [43]. The optimization problem was then solved using the BOBYQA optimizer implemented in the NLopt toolbox [44], [45]. To seed the BOBYQA optimizer, we used the closed-form solution provided by the C_AJ_ model.

## III Results

### A. Cost Estimation Accuracy

The accuracy of the parameterization of the proposed cost function was computed as the normalized root-mean-square error (RMSE) between the interpolated C_BM_ and a hold-out set of simulated biomechanical costs from *n* = 2100 arm postures (Fig. 3). The arm postures from the hold-out set were sampled as a 10 × 10 uniform grid for each DoF pair. For each pair, all other DoFs were kept static at their default values (as defined in the MSM). Given this analysis, the interpolated C_BM_ resulted in a normalized RMSE of 10.3312%.

**Fig. 3.**
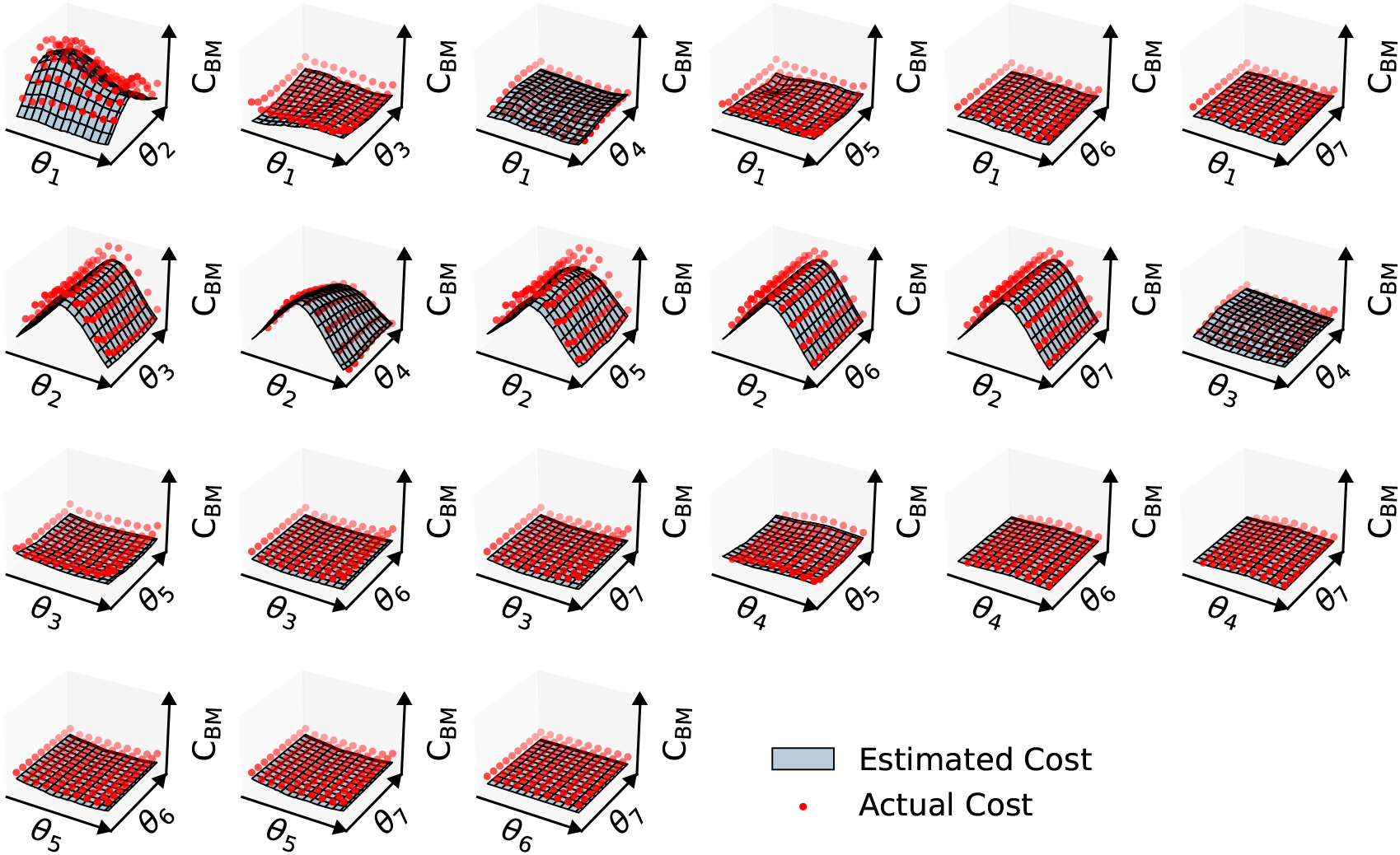
A visualization of the cost function, **C**_**BM**_. **C**_**BM**_ is computed as the interpolated surface in ℝ^**7**^ given the generalized forces generated by ***N* = 20000** inverse dynamics simulations of the MSM. **C**_**BM**_ is visualized as the 2D slice for each DoF pair (assuming all other DoFs are equal to their default value defined by the MSM). ***θ***_**1**_ refers to shoulder elevation plane, ***θ***_**2**_ refers to shoulder elevation magnitude, ***θ***_**3**_ refers to humeral rotation, ***θ***_**4**_ refers to elbow flexion, ***θ***_**5**_ refers to forearm rotation, ***θ***_**6**_ refers to wrist deviation, and ***θ***_**7**_ refers to wrist flexion. The interpolated **C**_**BM**_ resulted in a normalized RMSE of 10.3312 % when compared to biomechanical costs measured directly from simulation (red; ***n* = 2100**).

### B. Path Similarity

The performance of the proposed cost function was compared against the comparative models outlined in Table I. For the IL participants, the C_BM_ model generated optimal hand paths with an average alignment distance of 41.23 ± 21.80 (Fig. 4). In comparison, the comparative models generated optimal hands paths with the following average alignment distances: 202.15 ± 153.34 (C_HJ_), 338.98± 151.89 (C_AJ_), 252.80± 63.46 (C_TC_), and 235.01± 53.46 (C_EG_). The C_BM_ model performed significantly better than all comparative models (*p* < 0.001, Mann-Whitney U-test).

**Fig. 4.**
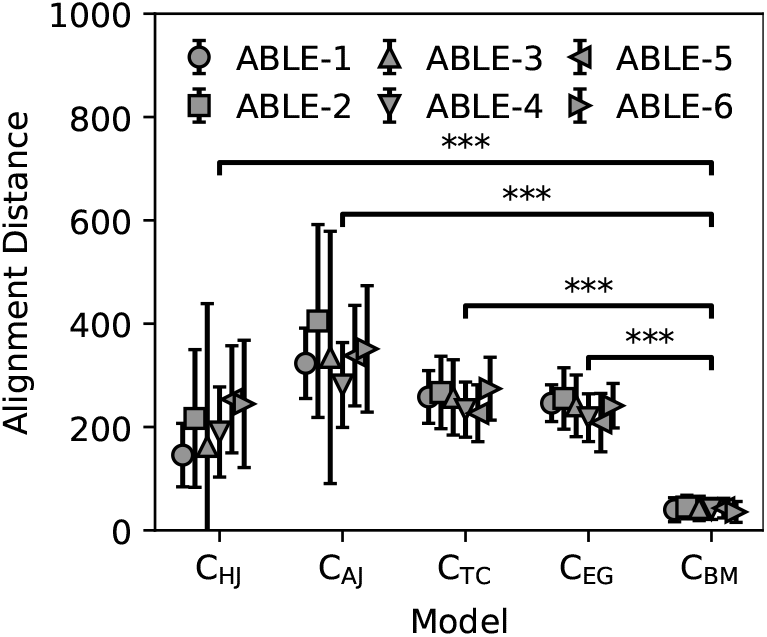
The alignment distance between measured reference hand paths and the optimal paths generated with various models of upper-limb motion. For IL participants, models generated optimal hand paths with the following mean alignment distances: 202.15 ± 153.34 (**C**_**HJ**_), 338.98 ± 151.89 (**C**_**AJ**_), 252.80 ± 63.46 (**C**_**TC**_), 235.01 ± 53.46 (**C**_**EG**_), and 41.23 ± 21.80 (**C**_**BM**_). Error bars denote 1 standard deviation. *** denotes ***p* < 0.001** (Mann-Whitney U-test).

For the ULL participant, the C_BM_ model outperformed the comparative models in both the *unloaded* and *loaded* experimental conditions (Fig. 5). For the ULL participant, we report performance of each model as the median alignment distance, followed by lower and upper quartiles: *M* (*Q*_*L*_, *Q*_*U*_ ). In the *unloaded* condition, the C_BM_ model generated optimal residual limb paths with an alignment distance distribution of 45.23 (29.24, 64.28). This is in comparison to the performance of the comparative models: 240.21 (162.74, 391.38) for C_HJ_, 949.39 (839.04, 1060.01) for C_AJ_, 301.19 (250.00, 386.25) for C_TC_, and 284.91 (250.79, 314.89) for C_EG_.

**Fig. 5.**
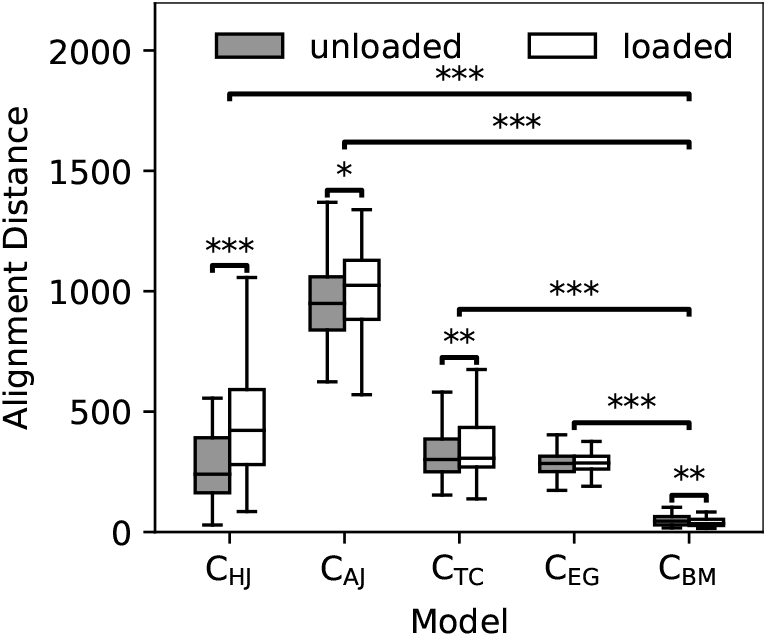
The alignment distance between measured reference residual limb paths and the optimal paths generated with various models of upper-limb motion. For the ULL participant, models generated paths with the following alignment distance distributions in the *unloaded* condition: 240.21 (162.74, 391.38) for **C**_**HJ**_, 949.39 (839.04, 1060.01) for **C**_**AJ**_, 301.19 (250.00, 386.25) for **C**_**TC**_, 284.91 (250.79, 314.89) for **C**_**EG**_, and 45.23 (29.24, 64.28) for **C**_**BM**_. For the *loaded* condition, models generated paths with the following alignment distance distributions: 422.05 (280.15, 591.64) for **C**_**HJ**_, 1024.58 (883.15, 1128.52) for **C**_**AJ**_, 306.24 (270.02, 434.27) for **C**_**TC**_, 286.59 (261.36, 314.75) for **C**_**EG**_, and 34.87 (27.00, 52.87) for **C**_**BM**_. * denotes ***p* < 0.05**, ** denotes ***p* < 0.01**, and *** denotes ***p* < 0.001** (paired Student’s t-test).

In the *loaded* condition, the C_BM_ model generated optimal residual limb paths with an alignment distance distribution of 34.87 (27.00, 52.87). The comparative models generated optimal paths with the following alignment distance distributions: 422.05 (280.15, 591.64) for C_HJ_, 1024.58 (883.15, 1128.52) for C_AJ_, 306.24 (270.02, 434.27) for C_TC_, and 286.59 (261.36, 314.75) for C_EG_. When compared to the *unloaded* experimental condition, every model (except C_EG_) showed a significant performance difference in the *loaded* condition (paired Student’s t-test). Of the models with load variant performance, C_HJ_, C_AJ_, and C_TC_ showed increased performance in the *unloaded* condition with *p* < 0.001, *p* < 0.05, and *p* < 0.01, respectively. In contrast, C_BM_ showed an increase in performance in the *loaded* condition, with *p* < 0.01.

## IV Discussion

In this manuscript, we propose an optimal upper-limb motor strategy generation framework that outperforms alternative methods in terms of geometric error (reported as alignment distance). The results of the 3D enpoint acquisition task show that the proposed cost function C_BM_ produces upper-limb motor paths with 1*/*5 of the alignment distance of the best-performing method previously proposed in the literature (C_HJ_): 41.23 vs 202.15. This small, mean alignment distance is accompanied by high model consistency, represented by 1*/*2 the standard deviation of the most consistent method previously proposed in the literature (C_EG_): 21.80 vs 41.23 (Fig. 4). These trends are present for both individuals with and without ULL as well as whether or not an individual with ULL is donning an upper-limb prosthesis (Fig. 5).

Furthermore, the C_BM_ model performs well across the entirety of an individual’s reaching volume (Fig. 6). While there are regions of low-error prediction shared among all models (e.g. 25^°^≤ *α* ≤ 45^°^, 45^°^≤ *β* ≤ 60^°^), the performance of the kinematic models, C_HJ_ and C_AJ_, varies greatly with the target hand position. This is in contrast to the C_TC_ and C_EG_ models, which present with less complex performance profiles, exhibiting a smaller range of alignment distances across the reaching volume. Still, the C_BM_ model performs the most consistently across the entire reaching volume, largely invariant to the desired hand position. These results are consistent with previous literature that has shown that optimal upper-limb motion is in part a function of the dynamics of the manipulator which varies nonlinearly with end-effector position [46]. Because C_TC_, C_EG_, and C_BM_ incorporate these dynamics inherently, their robust performance across the reaching volume is unsurprising.

**Fig. 6.**
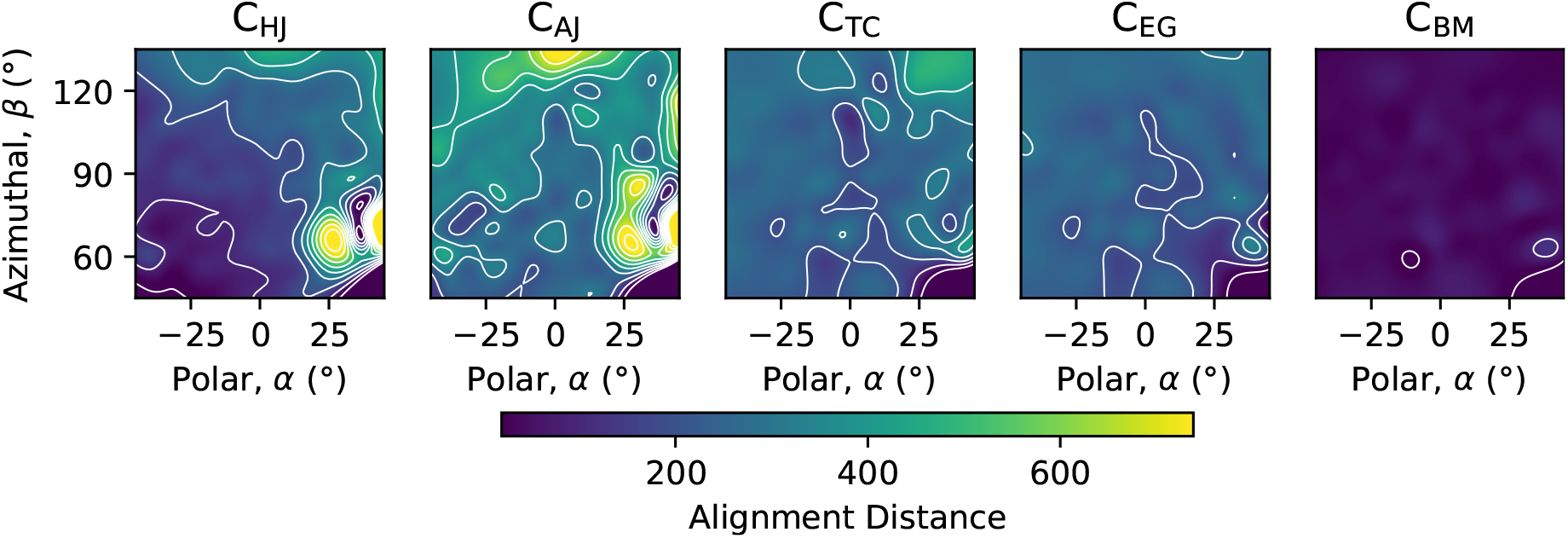
Average performance of various models as a function of endpoint position. Kinematic models, **C**_**HJ**_ and **C**_**AJ**_, present with the most complex performance profiles as the target hand position varies with polar and azimuthal angles. **C**_**TC**_ and **C**_**EG**_ present with less complex profiles, with a smaller range of alignment distances across the reaching volume. The performance of **C**_**BM**_ is largely invariant to the desired hand position. Contours of the performance space are highlighted in white.

**Fig. 7.**
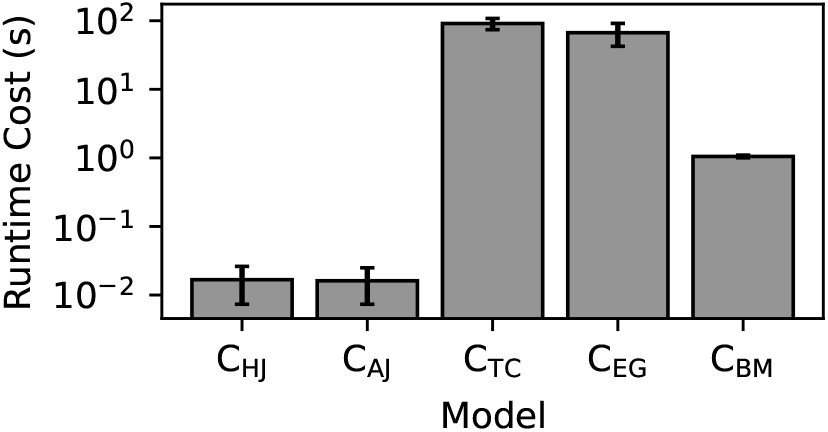
The time elapsed for each model to generate a single upper-limb motor path. Each model was used to generate 700 upper-limb paths on an i5-9400 processor (Intel, Santa Clara, CA). The models presented with the following run-time costs (in seconds): 0.0167 ***±*** 0.0094 (**C**_**HJ**_), 0.0161 ***±*** 0.0088 (**C**_**AJ**_), 91.1274 ***±*** 17.0331 (**C**_**TC**_), 66.9065 ***±*** 24.5498 (**C**_**EG**_), and 1.0491 ***±*** 0.0473 (**C**_**BM**_).

The accurate and robust performance of the C_BM_ model is accomplished with a per-path computational burden of 1.0491 ± 0.0473 s of runtime cost. While this is significantly greater than models with a closed-form solution, C_HJ_ (0.0167 ± 0.0094 s) and C_AJ_ (0.0161 ± 0.0088 s), this is an order of magnitude less than the models that require a numerical optimization, C_TC_ (91.1274 ± 17.0331 s) and C_AJ_ (66.9065 ± 24.5498 s). While 1 s of computation time is too slow for traditional real-time path planning, 1 s is brief enough for use in hybrid meta-planning control frameworks [47].

### A. Study Limitations

Our study was not without limitations. Most apparently, the experimental task was conducted with a limited ULL participant pool. With a single participant, the ULL results presented in this work act more as a case study. The results of this work can be strengthened by extending the experimental task to a larger population of ULL participants.

Another limitation of this study is the formulation of an upper-limb motor strategy as a *path*, rather than a *trajectory*. While a path is a continuous mapping from ***θ***_0_ to ***θ***_*f*_ , a trajectory is a path with a schedule. While the comparative methods generate upper-limb trajectories, and therefore provide additional temporal structure, the method proposed in this work generates upper-limb paths and is therefore purely geometric. This limits the analyses applicable to these upper-limb strategies, excluding temporal methods such as velocity profiles and motor phase durations. Of course, for control applications wherein temporal structure is required, the optimal paths generated from the proposed model can be transformed into trajectories using established methods [48].

Finally, our study is limited by the simplicity of the experimental task. While the 3D experimental task is more complex than the 2D tasks popularized in literature [9], [14], [15], [16], the unconstrained nature of the experiment allows flexibility in the optimal solution that is not representative of more functional prosthesis operation. For example, there is no constraint on the final hand or residual limb orientation, an important factor during object grasping. An extension of this work should replace the 3D target acquisition task with a clinical outcome measure in order to evoke motor strategies with more functional implications [49], [50].

### B. Future Directions

Future extensions of this work may explore a variety of improvements. For example, while this framework has performed well using a generalized MSM, there may be performance gains to be had by leveraging a subject-specific model of the upper-limb. This enhancement can leverage recent work that has developed software pipelines to automate the construction of personalized MSMs from medical imaging data [51]. This framework can also be extended to generate upper-limb motor strategies when (1) the environment contains obstacles and (2) upper-limb control is unreliable. To accomplish this, methods of obstacle avoidance and uncertainty-aware path planning can be applied to the C_BM_ cost space defined in this work (Fig. 3) [52], [53].

## V. Conclusion

This work outlines a framework for computing upper-limb motor strategies that outperforms alternative methods in terms of geometric error. This framework leverages musculoskeletal modeling to generate a cost space over which traditional path planning techniques can be applied to the biological limb. The proposed method showed decreased geometric error for predicted strategies in both individuals with and without an upper-limb amputation, both with and without an upper-limb prosthesis.

## Acknowledgment

The authors would like to thank the human subjects who participated in this study; Infinite Biomedical Technologies; the National Institutes of Health; and The Johns Hopkins University School of Medicine.

